# Untargeted Plasma Metabolomics in Canine Cognitive Dysfunction: The Naturally Occurring Alzheimer’s Disease Analog in Dogs

**DOI:** 10.1101/2025.06.20.660766

**Authors:** T. Melgarejo, S. Harrison, Y. Chang, M. Munoz, M. Kim, Y. Choi, J. Riveroll-Gonzalez, B. Natterson-Horowitz, A. Linde

**Author notes:** **CORRESPONDENCE** Tonatiuh Melgarejo, Department of Clinical Sciences, College of Veterinary Medicine, Western University of Health Sciences, 309 East 2nd Street, Pomona CA 91766, USA.

## Abstract

**Introduction:** Canine Cognitive Dysfunction (CCD) is an increasingly prevalent naturally occurring neurodegenerative condition in senescent dogs that share neuropathological and clinical features with human Alzheimer’s disease (AD). Metabolic profiling allows for identification of new candidates for AD biomarkers, diagnostics, and therapeutics. Despite its translational potential, plasma metabolomic profiling of dogs with CDD has not been previously characterized.

**Methods:** This case-control study analyzed plasma samples from ten client-owned geriatric dogs, including five with severe CCD and five age-matched, clinically healthy controls. Untargeted plasma metabolomics was performed using ultra-performance liquid chromatography–mass spectrometry (UPLC-MS). Multivariate and univariate statistical analyses identified significant metabolic differences between the groups. Metabolites were considered significant based on a variable importance in projection (VIP) score > 1.5, fold change (FC) > 2.0, and adjusted p-value < 0.05.

**Results:** Fifteen metabolites across seven chemical classes were significantly altered in CCD dogs compared to controls, including glycerophospholipids, steroid derivatives, indoles, and mitochondrial-related compounds. Notably, elevated lysophosphatidic acid (LPA 20:2/0:0) and reduced ubiquinone-2 levels suggest dysregulation in neuroinflammatory and oxidative stress pathways. Cholesterol exhibited the highest FC and VIP scores, further reinforcing its role in AD pathogenesis. Hierarchical clustering and pathway enrichment analyses supported distinct metabolic signatures in CCD that mirror those observed in human AD.

**Discussion:** This is the first untargeted plasma metabolomic profiling of dogs with CCD, revealing systemic metabolic disturbances that align with AD pathophysiology. Data was collected from senescent community-dwelling companion dogs, which enhances the study’s ecological and translational relevance. It supports the utility of CCD as an AD model and highlight candidate plasma biomarkers that warrant further investigation. Future longitudinal studies integrating metabolomics with neuroimaging, histopathology, and behavioral assessments are required to validate these findings and contribute to AD biomarker discovery and therapeutic development.

## 1. INTRODUCTION

Alzheimer’s disease (AD) represents the leading cause of dementia globally and is the fastest-growing epidemic ^1,2^. Deaths from dementia is considered underreported and some suggest AD may be the third leading cause of mortality in the US after cardiovascular disease and cancer ^3^. As a progressive neurodegenerative disorder marked by memory impairment, cognitive dysfunction, and behavioral changes, the pathology is characterized by accumulation of extracellular amyloid-beta plaques and intracellular neurofibrillary tangles composed of hyperphosphorylated tau protein. These changes are involved in driving synaptic dysfunction, neuronal loss, and progressive brain atrophy ^4^. The risk of developing dementia is significantly impacted by environmental factors, and a healthy lifestyle lowers the risk of AD substantially ^2^. However, despite decades of intensive research and major financial investments aimed at developing effective AD therapies, this pathology remains an incurable neurological disorder, which underscores the urgent need for innovative approaches to address this growing global health challenge ^5^. Rodent models have contributed to an increased understanding of AD pathobiology; however, these laboratory models do not develop AD-like pathology naturally and their translational utility is constrained by significant interspecies differences in brain architecture, immune function, and genetic composition. These disparities often result in findings that fail to accurately predict therapeutic outcomes in humans, thereby limiting the clinical relevance of rodent-based studies ^6^. The National Institute on Aging (NIA) has highlighted the critical need for alternative strategies to bridge this gap and pave a path towards successful AD drug development. Animal models that develop AD-like neurodegenerative conditions spontaneously are expected to provide more efficient and biologically relevant platforms for studying AD mechanisms and therapeutics. Still, the availability of naturally occurring animal models that recapitulate the complex pathology of AD with a high degree of fidelity is limited, which constitutes a roadblock for translational research ^7^. A few non-human animal species - including dog, degu, rhesus macaque, green monkey, and marmoset - are viewed as potential alternative models given their propensity for spontaneous AD-like neurodegenerative disease ^8,9^. The dog stands out as the only domesticated species, and companion dogs constitute a unique model system for more relevant and ethical AD research, because they live in human households with shared environmental exposures, unlike laboratory Beagle dogs.

Canine cognitive dysfunction (CCD) is an increasingly prevalent condition in geriatric dogs. Importantly, dog dementia is characterized as the analog of human AD ^10^. Aging dogs with CCD have been recognized as a unique and valuable animal model for AD research for over two decades due to the spontaneous isomorphic cellular changes that closely mirror human brain pathology and behavioral deficits ^11–13^. However, the dog is considered an expensive and technically challenging model, which has prevented widespread use in AD research. The relative void is primarily attributed to higher husbandry costs, long-term housing requirements, and the complexity of rigorous behavioral evaluations ^14^. Studies involving client-owned dogs have often yielded inconclusive results due to breakdown in owner compliance when clients euthanize dogs with CCD prematurely due to declining quality of life and increasing emotional disconnection. Despite the undeniable translational potential of the model, these factors have contributed to the underutilization of CCD as a naturally occurring animal model for AD research. The existing limitations of spontaneous animal models for preclinical research constitutes a barrier for development of effective disease-modifying therapies for AD ^7^. Consequently, a compelling rationale exists for re-evaluating the study of companion dogs with spontaneous CCD as a novel non-transgenic animal model for AD research.

Untargeted metabolomics is a powerful analytical approach that enables comprehensive profiling of small-molecule metabolites in biological samples, while offering insights into systemic physiological and pathological states. When applied to non-invasive biofluids such as plasma, this technique facilitates the identification of metabolic alterations associated with health and disease. The aim of this study was to characterize the untargeted plasma metabolomics profiles in a cohort of dogs with spontaneous CCD compared to controls. The research sought to advance comparative investigations into metabolic events implicated in the neurodegenerative processes associated with dementia in dogs of predicted future translational value for human AD research.

## 2. METHODS

### 2.1 Study design and aim

This is a single-center, case-control study aiming to comprehensively analyze plasma metabolomic profiles of client-owned, community-dwelling, companion dogs diagnosed with canine cognitive dysfunction (CCD) compared to age-matched clinically healthy controls.

### 2.2 Animals

A total of ten companion dogs older than 10 years of age were included in the study with prior client consent. The CCD group (n = 5) included dogs with a previously confirmed diagnosis of Canine Cognitive Dysfunction made by a licensed veterinarian based on clinical signs and a canine dementia scale (CADES) score above 44 points ^15^. The control group (n = 5) included age-matched clinically healthy companion dogs with a CADES score below 8 points. Exclusion criteria included significant hematologic or serum chemistry abnormalities, other significant co-morbidities, non-CCD neuropathology, psychopharmaceutical therapy, behavioral problems unrelated to CCD, and younger than 10 years of age.

### 2.3 Hematology and serum chemistry analysis

Blood (6 ml) was collected via cephalic or jugular venipuncture for a complete blood count (CBC), serum chemistry panel (SCP), and untargeted plasma metabolomics (UPM) analyses. The sample volume was divided evenly (2 ml per tube) into a red cap-yellow ring (serum clot activator), purple cap (EDTA), and green cap (lithium heparin) tubes, used for CBC, SCP, and UPM analyses respectively. The CBC and SCP analyses were performed at the WesternU Pet Health Center using a ProCyte Dx Hematology Analyzer (CBC), and Catalyst One Chemistry Analyzer (SCP), both from IDEXX Laboratories (Westbrook, ME). Data was analyzed using MS Excel for Mac (Ver. 16.97.2) for descriptive and inferential statistics (t-test, alpha level of 0.05).

### 2.4 Sample preparation for untargeted plasma metabolomics

Whole blood samples for UPM (2 ml in lithium heparin tubes) were centrifuged at 3,000 rpm x 10 min and 1 ml of plasma was collected from each sample, stored at -80°C and batched prior to overnight shipping on dry ice to Creative Proteomics (CP) (Shirley, NY) for untargeted plasma metabolomics analysis. The following protocol was provided by CP. Briefly, samples were thawed on ice and 60 μL of each sample was subsequently transferred into a tube containing 1.5 mL of chloroform:methanol (2:1, v/v) and 0.5 mL of ultrapure water. The mixture was homogenized for 90 sec., vortexed for 1 min, and sonicated for 30 min at 4°C. Samples were then centrifuged at 3,000 rpm for 10 min at 4°C, and the lower phase transferred to a new tube and dried under nitrogen gas. The dried extract was resuspended in 200 μL isopropyl alcohol:methanol (1:1, v/v), followed by addition of 5 μL LPC (12:0) as an internal standard. After centrifugation at 12,000 rpm for 10 min at 4°C, the supernatant was collected for liquid chromatography–mass spectrometry (LC-MS) analysis. Quality control (QC) samples were prepared by pooling equal amounts of extract from each sample, following the same sample preparation procedure. QC samples were included in the analysis to assess the stability and reproducibility of the method.

### 2.5 Ultra-performance liquid chromatography-mass spectrometry (UPLC-MS)

Metabolite separation was performed by CP using ultra-performance liquid chromatography (UPLC) coupled with a Q Exactive mass spectrometer (Thermo Fisher Scientific, Waltham, MA). Chromatographic separation was achieved using an ACQUITY UPLC BEH C18 column (100 × 2.1 mm, 1.7 μm). The mobile phase consisted of solvent A (60% ACN+40% H2O+10 mM HCOONH4) and solvent B (10% ACN+90% isopropyl alcohol+10 mM HCOONH4). A gradient elution program was applied, starting with 30% B from 0 to 1.0 min, increasing to 100% B from 1.0 to 10.5 min, maintaining at 100% B from 10.5 to 12.5 min, and returning to 30% B from 12.5 to 12.51 min, where it remained until 16 min. The flow rate was set at 0.3 mL/min, with the column temperature maintained at 40°C and the sample manager temperature at 4°C. For mass spectrometry, both electrospray ionization positive (ESI+) and negative (ESI−) modes were utilized. In ESI+ mode, the heater temperature was 300°C, sheath gas flow rate was set at 45 arbitrary units (arb), auxiliary gas flow rate at 15 arb, sweep gas flow rate at 1 arb, spray voltage at 3.0 kV, capillary temperature at 350°C, and S-Lens RF level at 30%. In ESI− mode, similar parameters were applied, except for the spray voltage, which was set at 3.2 kV, and the S-Lens RF level, which was set at 60%.

### 2.6 Statistical analysis and metabolite identification

Raw data was acquired by CP and aligned using Lipid Search software (Thermo Fisher Scientific, Waltham, MA), with ion signals from both ESI+ and ESI− modes merged. Principal Component Analysis (PCA) was used as an unsupervised method to visualize data clustering and identify potential outliers. To further distinguish metabolic differences between groups, Partial Least Squares Discriminant Analysis (PLS-DA) was performed using Python’s PLSRegression, while Orthogonal Partial Least Squares Discriminant Analysis (OPLS-DA) was conducted using Python’s pyopls. Data visualization was implemented using Matplotlib. The False Discovery Rate (FDR) correction, specifically the Benjamini-Hochberg method, was applied to control for multiple testing and to balance sensitivity and specificity.

Metabolites were considered significant if they met the following criteria: a Variable Importance in Projection (VIP) score greater than 1.5, a Fold Change (FC) greater than 2.0, and a Benjamini-Hochberg-adjusted p-value below 0.05. Hierarchical clustering analysis (HCA) was performed using the complete linkage algorithm in Cluster 3.0 (Stanford University, Stanford, CA), with visualization carried out using Seaborn’s clustermap function. Metabolite identification was conducted using the Human Metabolome Database (HMDB) and confirmed by MS/MS fragment analysis. Pathway analysis was performed using KEGG and MetaboAnalyst 5.0 to determine the metabolic pathways most affected in CCD.

## 3. RESULTS

### 3.1 Study subjects and groups

Dogs enrolled in the study were client-owned, altered (female spayed, male neutered) with a median age of 13 years (range 10 - 16 yrs.) and 40% were male. Breeds included Australian shepherd, boxer, dachshund, German shorthaired pointer, Labrador retriever mix (2), and Chihuahua (4). Signalment and health status of all study subjects is included in Table S1. For the CCD group, dogs were confirmed to have severe cognitive impairment based on clinical signs and a CADES score ≥ 45. Control dogs had a CADES score ≤ 7 (Table S2).

### 3.2 Complete blood count and serum chemistry panel

Comprehensive hematological (CBC) and biochemical (SCP) profiling revealed neither clinically relevant abnormalities in either group nor statistically significant differences between the groups (Table S3).

### 3.3 Plasma untargeted metabolomics

#### 3.3.1 Quality control assessment

To ensure the stability of the LC-MS system, QC samples were analyzed at regular intervals in both positive and negative ion modes. The relative standard deviation (RSD) distribution, shown in Figure S1, indicated that most RSD values were below 30%, confirming the robustness of the analytical method.

#### 3.3.2 Multivariate statistical analysis

Normalization was performed to ensure all samples were within a 95% confidence interval, confirming system stability. PCA analysis revealed clear clustering patterns, reflecting distinct metabolic variations between groups. Further analysis using PLS-DA demonstrated significant separation between the CCD and clinically healthy control (CH) group (Figure 1.A). Similarly, OPLS-DA score plots (Figure 1.B) confirmed the metabolic differences between the two groups.

**Figure 1.**
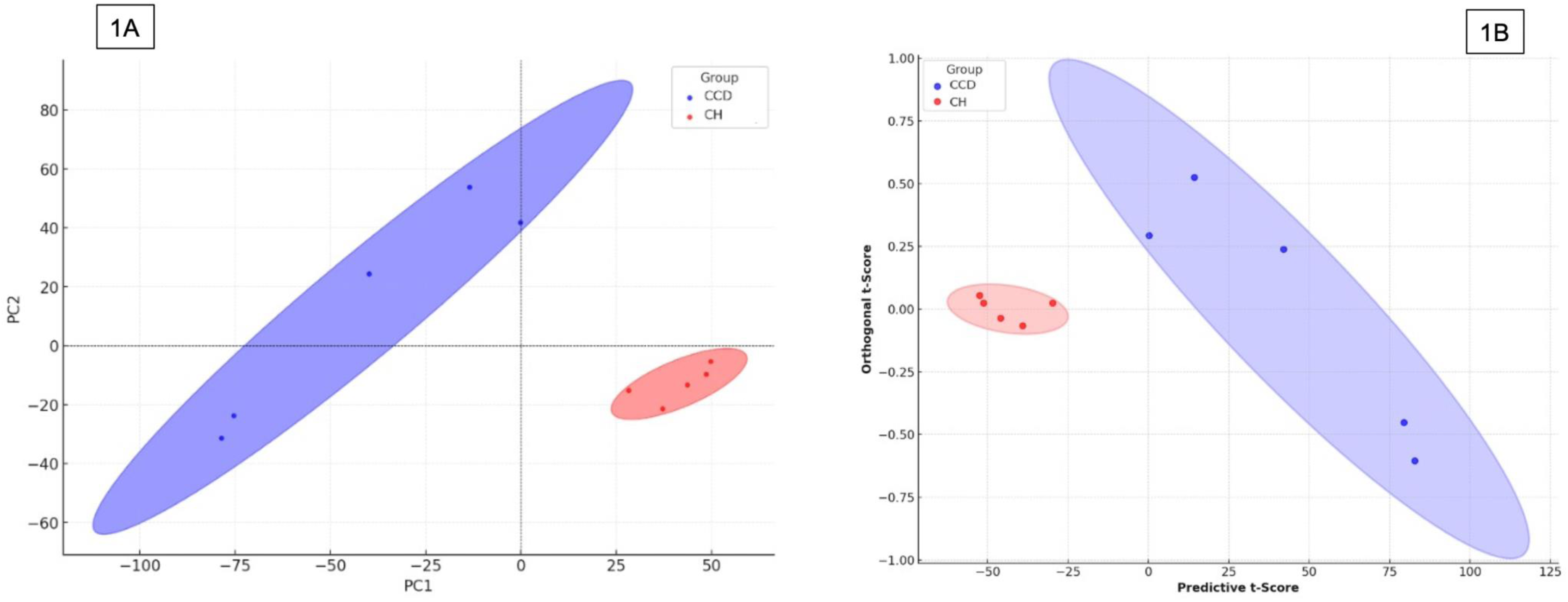
A) Partial least squares discriminant analysis (PLS-DA) score plot demonstrating clear separation between dogs with canine cognitive dysfunction (CCD) (blue) and clinically healthy controls (CH) (red) groups. Each point represents an individual sample, and the ellipses reflect 95% confidence intervals for each group. The first principal component (PC1) accounts for 74.39% of the total variance, while the second principal component (PC2) explains 25.61%, highlighting a distinct clustering pattern between the two groups. B) Orthogonal partial least squares discriminant analysis (OPLS-DA) score plot illustrating group separation between CCD (blue) and CH (red). Each point represents an individual sample, and the shaded ellipses denote the 95% confidence intervals for each group. The predictive t-score (horizontal axis) explains 39% of the variation, while the orthogonal t-score (vertical axis) accounts for 25.6%, indicating distinct clustering between the two groups based on their multivariate profiles.

#### 3.3.3 Single variable analysis

Metabolites were selected as significant if they met the criteria of VIP > 1.5, FC > 2.0, and an adjusted p-value below 0.05 (Table 1). Using these criteria, fifteen AD-associated metabolites were identified including glycerophospholipids, indoles, polyprenylbenzoquinones, pyridines, saturated hydrocarbons, and steroid derivatives. Cholesterol, although not meeting the adjusted p-value threshold (*p* = 0.11), was included as a biologically relevant metabolite due to having the highest FC and VIP among all significant plasma metabolites combined with existing evidence linking elevated cholesterol levels to amyloid-beta accumulation, neuroinflammation, and vascular contributions to cognitive impairment ^16^. The volcano plot in Figure 2 highlights key metabolic differences between the groups (CCD vs CH).

**Table 1.**
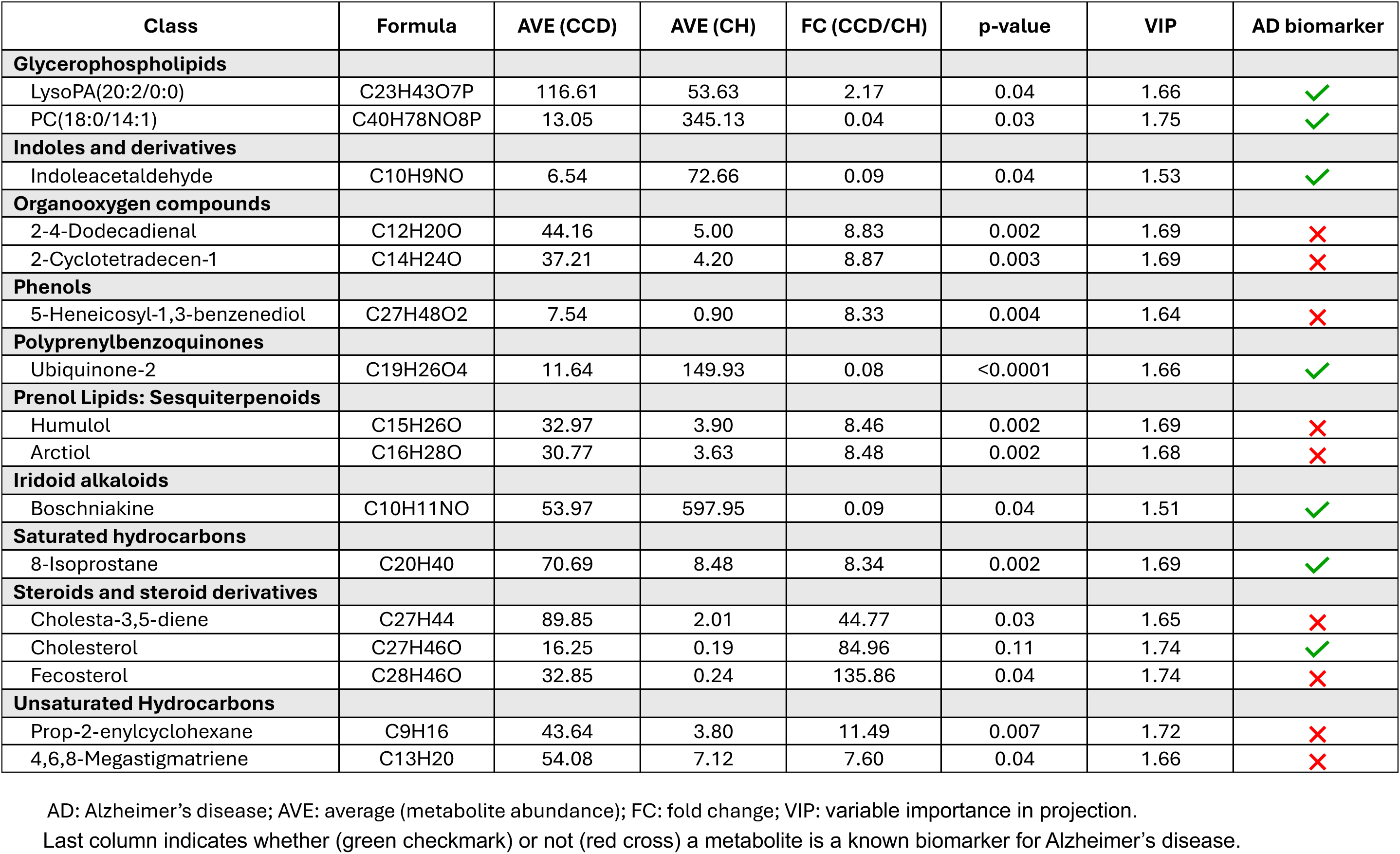
Select plasma metabolites in dogs with canine cognitive dysfunction (CCD) vs clinically healthy controls (CH)

**Figure 2.**
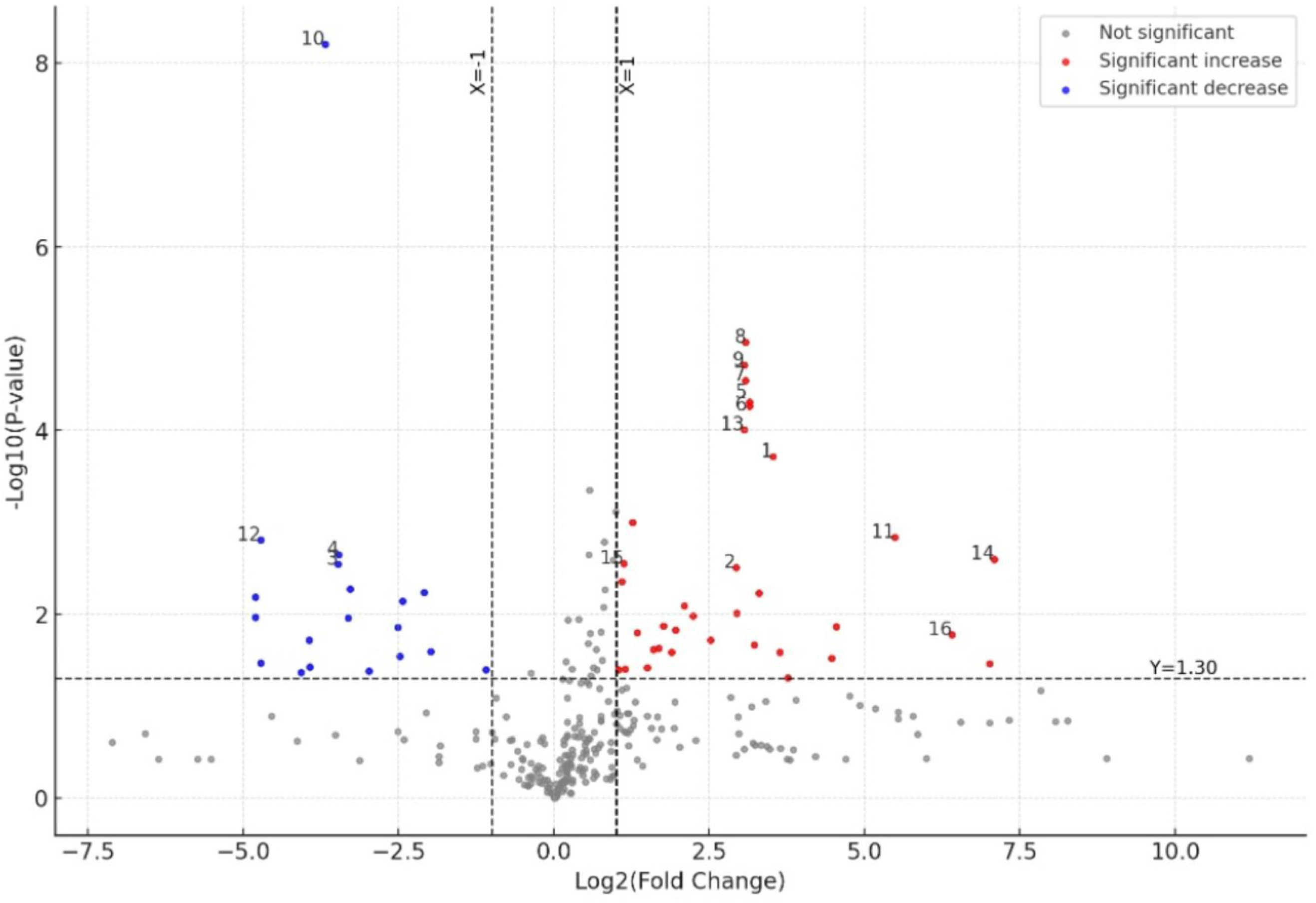
Volcano plot illustrating differential metabolite expression between the canine cognitive dysfunction (CCD) group vs the control group. The x-axis represents the log₂(fold change), and the y-axis shows the –log₁₀(p-value). The horizontal dashed line corresponds to the significance threshold (FDR-adjusted p-value < 0.05; –log₁₀(p) ≈ 1.30), and the vertical dashed lines indicate the fold change cutoffs (log₂FC = ±1). Metabolites significantly upregulated in the CCD group are shown in red, while those significantly downregulated are shown in blue. Metabolites that did not meet the significance threshold are shown in gray. Select metabolites of specific interest are annotated by number as follows: 1: Prop-2-enylcyclohexane; 2: 4,6,8-Megastigmatriene; 3: Indoleacetaldehyde; 4: Boschniakine; 5: 2-4-Dodecadienal; 6: 2-Cyclotetradecen-1; 7: Humulol; 8: Arctiol; 9: 8-Isoprostane; 10: Ubiquinone-2; 11: Cholesta-3,5-diene; 12: PC(18:0/14:1); 13: 5-Heneicosyl-1,3-benzenediol; 14: Fecosterol; 15: LysoPA(20:2/0:0); 16: Cholesterol

#### 3.3.4 Cluster analysis

To identify trends in metabolite variation, mean metabolite content from the groups was used to calculate metabolite ratios. After log transformation and normalization, hierarchical clustering analysis (HCA) was performed. The heatmap in Figure 3 illustrates the distribution of significant metabolites (plus cholesterol) in dogs with CCD vs CH (controls), where red represents increased levels and green indicates decreased levels relative to the median.

**Figure 3.**
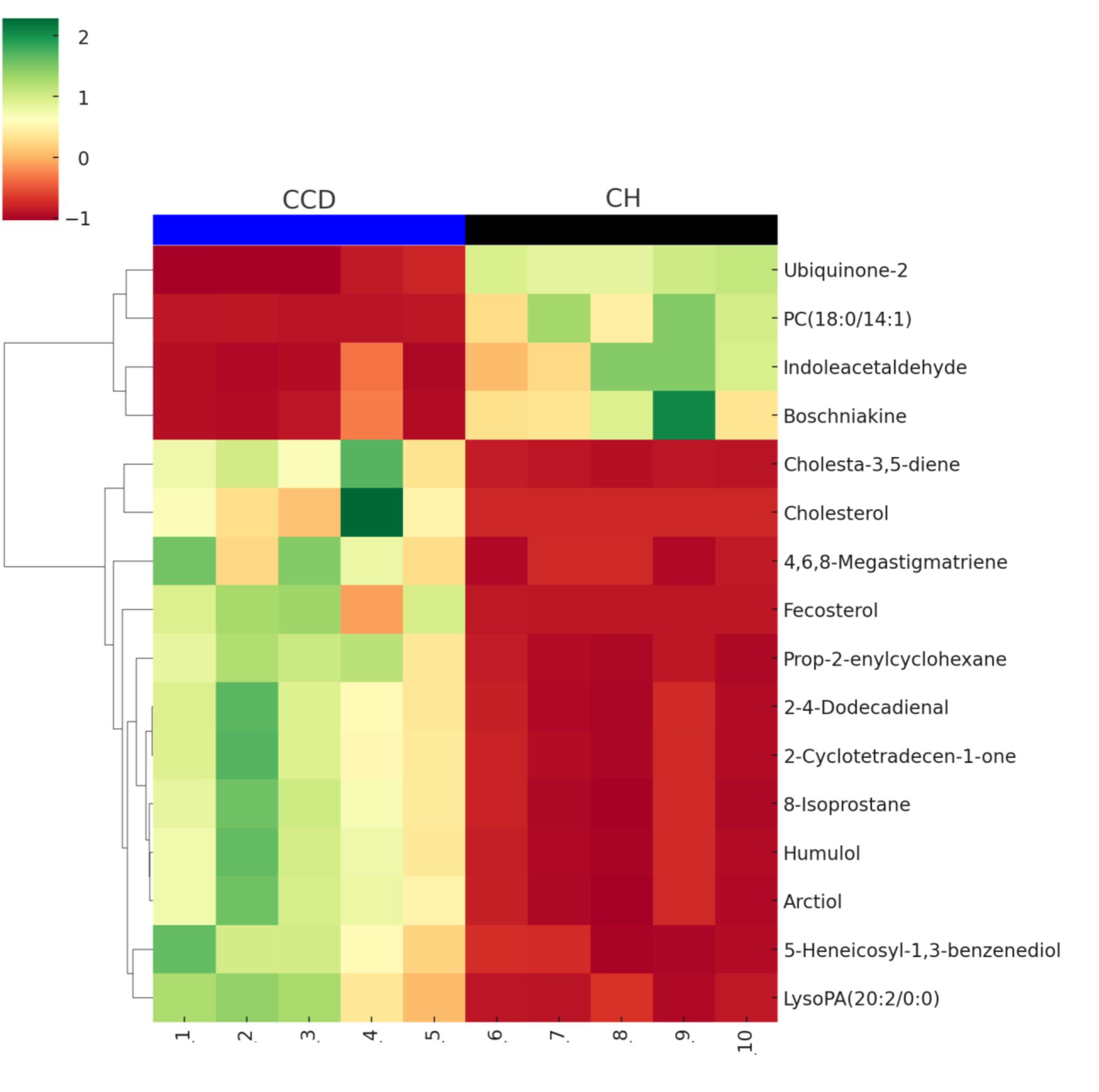
Heatmap illustrating the distribution of select metabolites across the canine cognitive dysfunction (CCD) vs clinically healthy control (CH) groups. Each row represents a metabolite, and each column corresponds to an individual sample (CCD: 1-5, CH: 6-10). Color intensity reflects relative abundance, with green indicating increased levels and red indicating decreased levels relative to the median for each metabolite. Hierarchical clustering was applied to group metabolites based on similarity in expression patterns.

#### 3.3.5 Metabolite correlation network

A metabolome view (Figure 4) was generated based on pathway enrichment and topology analysis. Each node in the network represents a metabolite set, with its size reflecting fold enrichment and its color intensity corresponding to statistical significance. Metabolite sets were considered connected when they shared over 25% of their combined metabolites, with the color gradient ranging from pink to red, indicating varying levels of significance.

**Figure 4.**
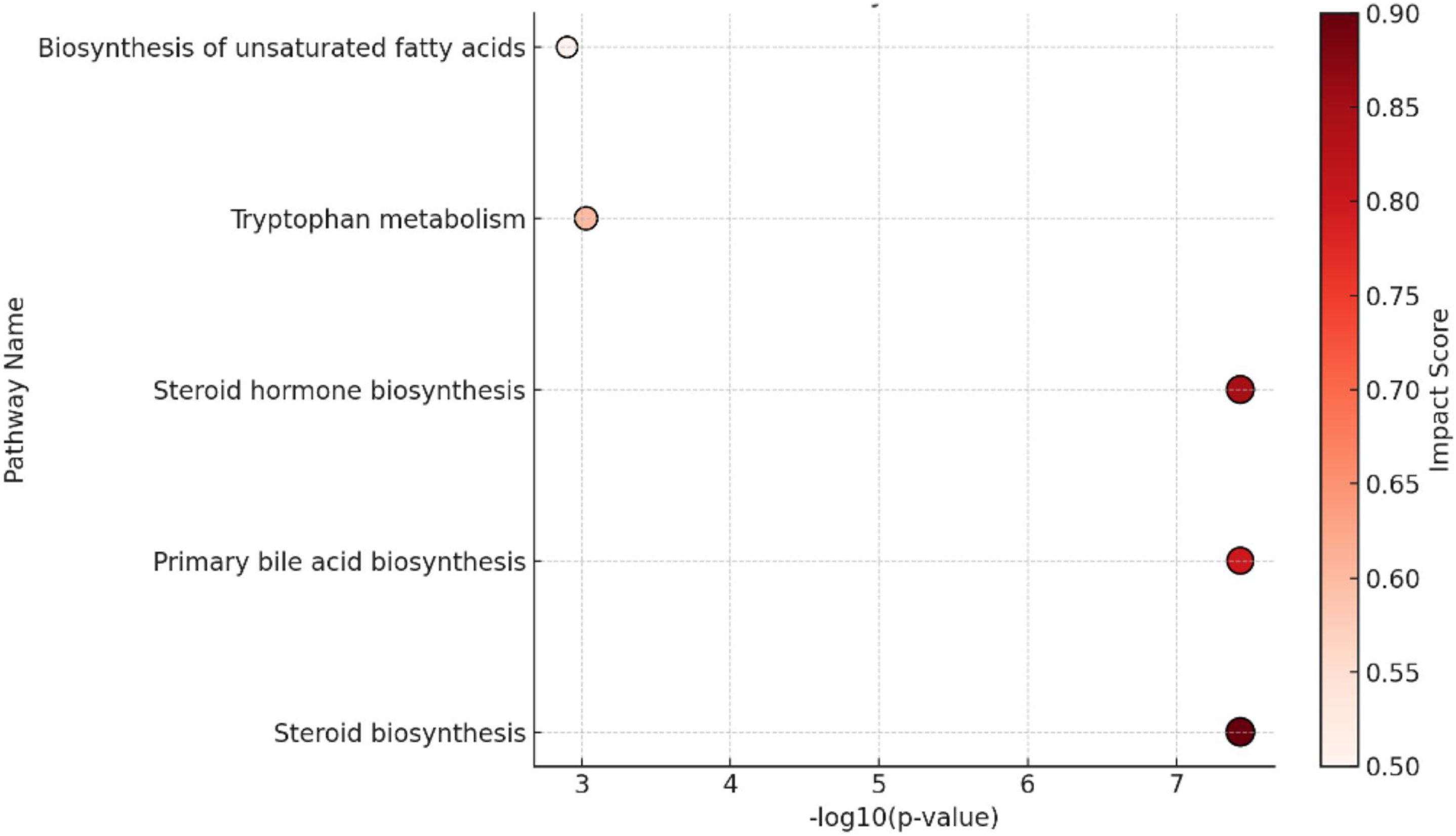
Pathway enrichment and topology analysis of perturbed metabolites between the canine cognitive dysfunction (CCD) and the clinically healthy control (CH) groups. The dot plot displays the top enriched metabolic pathways, with the x-axis representing pathway significance (–log₁₀(p-value)) and the color intensity indicating the pathway impact score, based on pathway topology analysis. Larger and darker red circles represent pathways with higher impact scores and greater biological relevance. Steroid biosynthesis, primary bile acid biosynthesis, and steroid hormone biosynthesis were among the most significantly enriched and impactful pathways.

## 4. DISCUSSION

This study provides a comprehensive analysis of plasma metabolomic profiles in companion dogs diagnosed with severe CCD compared to age-matched clinically healthy controls. Utilizing an untargeted metabolomic approach, our investigation revealed significant alterations in multiple plasma metabolite classes in the CCD group that mirror those associated with AD in people, including glycerophospholipids, indoles, polyprenylbenzoquinones, iridoid alkaloids, saturated hydrocarbons, and steroid derivatives. Our data revealed significant differences in plasma glycerophospholipid concentrations between the groups (CCD vs CH). Specifically, there was a marked increase in lysophosphatidic acid (LPA) 20:2/0:0 in CCD dogs compared to controls. LPA 20:2/0:0, a bioactive lipid, plays a crucial role in cellular signaling processes. LPAs (including LPA 20:2/0:0) have been implicated in several key pathological mechanisms in AD. Notably, LPAs activate microglia and astrocytes, leading to the release of pro-inflammatory cytokines, which significantly contribute to the progression of AD through neuroinflammation ^17^. LPAs enhance amyloid-beta (Aβ) production by upregulating beta-secretase expression, which is crucial for formation of amyloid plaques, a hallmark of both AD and CCD ^12,17^. Additionally, LPAs exacerbate neurodegeneration by compromising the integrity of the blood-brain barrier, allowing harmful substances to enter the brain. LPAs are moreover involved in phosphorylation of tau proteins, leading to formation of neurofibrillary tangles, another key feature of AD pathology ^17^. We detected elevated LPA 20:2/0:0 plasma concentration in the CCD group, which is relevant as recent studies have shown that aged canines with CCD exhibit phosphorylated tau at threonine 217 ^18^. The study also showed that PC 18:0/14:1 was significantly decreased in the CCD group, which is a plasma glycerophospholipid essential for maintaining cellular membrane integrity and fluidity. This finding runs parallel to metabolomics data in people with AD. The reduction in PC 18:0/14:1 appears to have similar biological effects to the increase of LPA 20:2/0:0 in humans with AD, namely, enhancing neuroinflammation ^19^, amyloid-beta (Aβ) aggregation ^20^, and exacerbation of tau hyperphosphorylation ^20^. Therefore, plasma glycerophospholipid alterations in dogs with CCD seem to be a crucial component of the neurodegenerative processes in the canine brain and should be considered a comparative biomedical tool to further enhance our understanding of AD pathophysiology.

In this study, the plasma concentration of ubiquinone-2, a key component of the mitochondrial electron transport chain, was significantly reduced in the CCD group compared to controls. The role of the coenzyme-Q class molecules (including ubiquinone-2 and CoQ10) has been investigated in the pathogenesis of AD, primarily focusing on their involvement in abnormal mitochondrial metabolism and oxidative stress ^21^. The decrease in ubiquinone-2 in dogs with CCD most likely reflects mitochondrial dysfunction, a hallmark of AD pathogenesis ^22,23^. Oxidative stress is one of the primary mechanisms involved in the pathogenesis of AD and other neurodegenerative disorders. Several studies have focused on analyzing the CoQ10 concentration in various tissues in AD patients and individuals with other dementia syndromes. These studies have also explored the potential therapeutic role of CoQ10 in AD treatment. The potential therapeutic effects of CoQ10 have been demonstrated in experimental models of AD, showing significant neuroprotective properties ^24^. In comparison, a recent systematic review and meta-analysis of studies measuring tissue CoQ10 levels in people with dementia vs controls revealed that AD patients have similar serum/plasma CoQ10 levels compared to controls ^24^. Studies in humans with AD and mild cognitive impairment (MCI) have yielded inconclusive results ^25^, while studies in rodent models shows a promising role of CoQ10 in improving memory ^26^. As such, this opens a door to study the use of CoQ10 in dogs with dementia to define potential translational benefits for people with AD and MCI.

Interestingly, we demonstrated a substantially higher plasma concentration of indole-acetaldehyde in the clinically healthy controls vs the CCD group. Indole derivatives, including indoleacetaldehyde, have been investigated for their ability to inhibit monoamine oxidase (MAO) enzymes as well as cholinesterase enzymes, which are involved in the breakdown of neurotransmitters. Inhibition of these enzymes can help in reducing oxidative stress and neuroinflammation ^27^. Studies have demonstrated that certain indole derivatives, including those with indoleacetaldehyde moieties, exhibit significant neuroprotective effects against neurotoxic insults in cellular models. These compounds show improved chemical stability that make them promising candidates for AD therapeutics development ^28^. Indole derivatives are primarily produced in the human body through metabolism of tryptophan by intestinal microorganisms ^29^. The gut microbiota plays a crucial role in converting tryptophan into indole derivatives that are released in the plasma where they can exert a variety of biological effects. The reduction in indoleacetaldehyde plasma concentration in the CCD group points to alterations in tryptophan metabolism, which has also been associated with neuroinflammation and neurodegeneration in AD ^30^. Since indole derivatives can influence gut-brain axis signaling, their dysregulation in CCD may reflect similar pathways operative in AD, thus emphasizing the translational relevance.

We found that the plasma concentration of Boschniakine, an iridoid alkaloid from the plant *Boschniakia rossica* (BR), was ten times higher in controls compared to the CCD group. BR contains more than 100 chemical constituents, most prominently boschnaside, boschniakine, 7-deoxy 8-epiloganic acid, and (4R)-4-hydroxymethyl-boschnialactone ^31^. These compounds possess a wide spectrum of pharmacological effects, including antineoplastic, antioxidant, and anti-inflammatory properties ^31^. BR has been used in China to treat age-related diseases, including dementia, for centuries, but the mechanism of action and potential therapeutic metabolic targets remain unclear. Recent pathological and behavioral studies in a rodent model of AD demonstrated that BR can significantly increase the number of nerve cells in the hippocampus and improve the learning and memory abilities of rats with AD-like disease ^32^. These results suggest that BR may improve neuronal health by modifying the course of the inflammatory response in AD. None of the commercial dog food sold on the US market incorporate unique botanical extracts like BR, which is an obligate parasitic plant that lacks chlorophyll. BR is native to the Northern Hemisphere and does not grow south of the Alaska Panhandle ^33^. We speculate that the source of BR may be derived from an association with plant ingredients commonly used in dog food manufacturing, such as blueberry and white willow. The data from the rodent model of AD, combined with our findings that clinically healthy dogs exhibit significantly elevated BR concentrations compared to dogs with CCD, establish a theoretical foundation that would encourage further investigation.

F2-isoprostanes, including the isomer 8-isoprostane, are products of non-enzymatic peroxidation of arachidonic acid, a polyunsaturated fatty acid in cell membranes. Growing evidence links oxidative stress and lipid peroxidation to the pathogenesis of neurodegenerative diseases such as AD ^34^. Elevated levels of F2-isoprostanes have been observed in neurons, cerebrospinal fluid, plasma, and urine of individuals with AD and MCI, indicating oxidative damage in the brain ^35,36^. In this study, the CCD group had a significantly higher plasma 8-isoprostane level, suggesting increased brain oxidative stress. While the causal relationship between F2-isoprostane elevation and AD remains under investigation ^37^, these findings have translational relevance. Companion dogs represent a unique spontaneous animal model that mirrors the pathways implicated in human AD, where oxidative damage is a key contributor to neurodegeneration ^38^.

Cholesterol regulation in the brain involves complex biochemical pathways. Steroid derivatives, like cholesterol, are essential for neural health, particularly in synapse formation and signal transmission. Disruptions in cholesterol metabolism are linked to neurodegeneration and are considered key contributors to AD pathogenesis, especially in late-onset cases where major genetic risk factors are tied to cholesterol regulation ^39,40^. This underscores a need for further investigation into cholesterol’s role in neurodegenerative diseases. In our study, the CCD group showed the second highest plasma cholesterol FC and VIP among the select plasma metabolites (Table 1). While the mechanisms remain unclear, cholesterol is known to influence amyloid precursor protein processing and Aβ accumulation in the human brain ^39^, which suggests that similar pathways may exist in dogs with CCD. In fact, the Aβ peptide family includes AD-associated peptides with identical amino acid sequences between dogs and humans ^12^. Notably, Aβ42 deposition in three brain areas correlated strongly with canine cognitive dysfunction scale (CCDS) scores ^41^, supporting the possible link between altered cholesterol metabolism and Aβ42 pathology in this species.

Nine (9) other circulating metabolites, though not directly linked to CCD or AD, showed significant plasma concentration changes, including organooxygen compounds (2), phenols (1), prenol lipids: sesquiterpenoids (2), steroids and steroid derivatives (2) and unsaturated hydrocarbons (2) (Table 1). These shifts suggest broader metabolic disruptions affecting energy balance, membrane function, and signaling, reinforcing the notion that the complex biochemical disturbance in CCD resembles that in patients with AD.

### 4.1 Strengths and limitations

Unlike laboratory animals with experimentally induced disease, companion dogs co-exist with people and share their environmental exposures. As such, the study of aging client-owned dogs with naturally occurring neurodegenerative disease boosts translational relevance. The relatively small sample size limits the generalizability of the results and dictates cautious interpretation. However, the pronounced metabolic alterations in dogs with severe cognitive impairment warrant further investigation.

### 4.2 Conclusion

Metabolomic profiling of aged dogs with CCD reveals systemic metabolic disturbances, particularly in lipid metabolism, mitochondrial function, and oxidative stress that closely resemble those observed in humans with AD. These findings reinforce the translational value of client-owned companion dogs as a spontaneous animal model for AD research. Unlike traditional laboratory models that involves purpose bred Beagles, companion dogs exhibit great genetic diversity and live in home environments that reflect real-world exposures, which makes them uniquely suited for studying the complex interplay between environmental exposures, aging, and neurodegeneration. Prospect studies with larger, longitudinal cohorts are essential to validate these candidate biomarkers and clarify their temporal association with cognitive decline. Integrating metabolomic data with neuroimaging, histopathology, and behavioral assessments will be critical to strengthening the mechanistic understanding and translational potential of this model in AD research. Continued exploration of these pathways holds promise for advancing translational efforts of relevance to effective management through novel disease-modifying therapies for Alzheimer’s disease. Precedent for the proposed strategy is demonstrated by the success of the National Cancer Institute’s Comparative Oncology Program and the Comparative Oncology Trials Consortium, which is dedicated to the study of naturally occurring cancers in companion dogs as a model system for human disease ^42^. Senescent companion dogs with CCD could similarly support comparative gerontology and AD focused research within a National Institute on Aging framework.

## AUTHORS CONTRIBUTIONS

Study concept and design: T.M, A.L; data acquisition: T.M, A.L, S.H, Y.C, M.M, J.R.G; data analysis and interpretation: T.M, Y.C, S.H, A.L; drafting of the manuscript: T.M, S.H, M.K, A.L; critical revision of the manuscript for important intellectual content: T.M, A.L, B.N.H, S.H, Y.C, M.M, M.K, Y.C, J.R.G.

## ACKNOWLEDGEMENTS

We thank all clients who volunteered their companion dogs for enrollment in this study. We also thank clinical staff and other colleagues in the College of Veterinary Medicine who helped facilitate this research.

## DATA AVAILABILITY STATEMENT

All data is presented either in the manuscript or the supplementary material.

## IACUC AND CLIENT CONSENT STATEMENT

Approved by the Western University of Health Sciences Institutioal Animal Care and Use Committee (protocol no. R23IACUC010). Client consent was obtained for all companion dogs enrolled in the study.

## Figure legends

**Figure S1.**
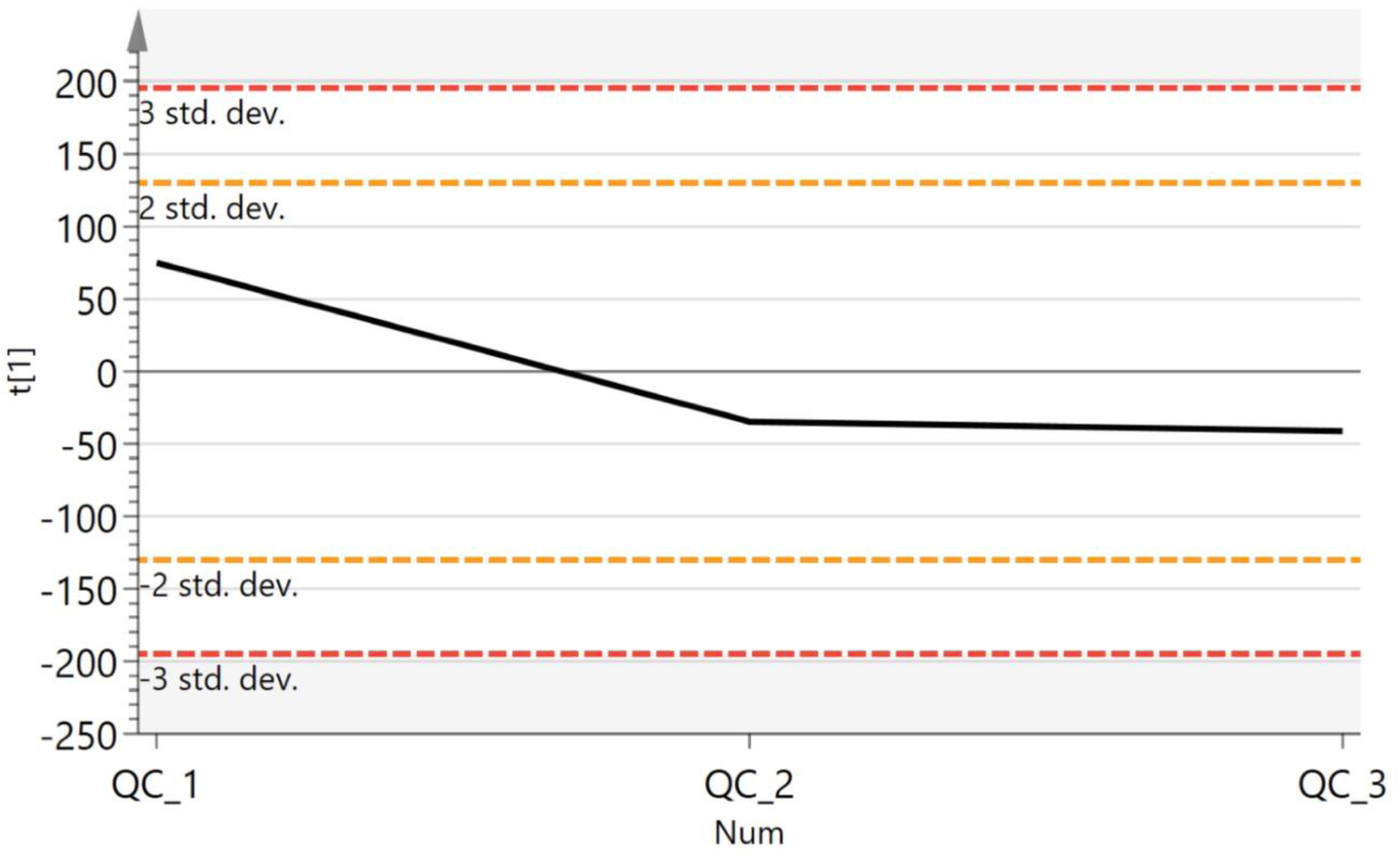
Quality control assessment of LC-MS system performance. To monitor analytical stability throughout the metabolomics run, pooled quality control (QC) samples were injected at regular intervals in both positive and negative ionization modes. The distribution of relative standard deviation (RSD) values across detected features is shown. Most features exhibited RSDs below 30%, indicating high reproducibility and robustness of the LC-MS platform used in this study.

## RESEARCH IN CONTEXT

### Systematic Review

The literature review was conducted using PubMed and Google Scholar. Comparative studies in Alzheimer’s disease research have overwhelmingly focused on rodent models, with a smaller percentage of studies targeting non-human primates, zebra fish, and invertebrates. These model systems have all shown limited translational value, documented by the lack of effective therapies for AD. Client-owned companion dogs are recognized as a valuable naturally occurring animal model for human AD due to dogs’ spontaneous aging and cognitive decline, shared environments and lifestyles, biological similarities between dementia in the two species, as well as validated cognitive assessment tools for dogs. The logistics required for large scale clinical research involving companion dogs, including those with dementia, is complex, as demonstrated by the Dog Aging Project (https://dogagingproject.org/). Metabolomics analysis provides unique insight on system effects of neurodegenerative processes; however, no peer-reviewed studies have characterized plasma metabolomic profiles in dogs with Canine Cognitive Dysfunction (CCD). <u>Interpretation</u>: This study reveals distinct plasma metabolomic alterations in companion dogs with CCD that mirror human AD, particularly in lipid metabolism, mitochondrial function, and oxidative stress. The use of naturally aging, client-owned dogs enhance translational relevance, offering a valuable model for investigating spontaneous neurodegeneration in real-world environmental contexts.

### Future Directions

Research should focus on validating these candidate biomarkers in larger, longitudinal cohorts and determining their temporal relationship with cognitive decline. Integrating metabolomics with neuroimaging, histopathology, and behavioral data will be essential to elucidate mechanisms and enhance the translational utility of this promising, yet largely unexplored, canine model in comparative AD research.

## SUPPLEMENTARY TABLES

**Table S1.** Signalment of canine study participants.

| Dog ID | Age (yr) | Dog Breed(s) | Gender | Weight (kg) | CADES score | Group |
| --- | --- | --- | --- | --- | --- | --- |
| 1 | 11 | Dachshund | MN | 9.6 | 79 | CCD |
| 2 | 16 | Lab mix | MN | 18.8 | 64 | CCD |
| 3 | 14 | Chihuahua mix | MN | 3.7 | 86 | CCD |
| 4 | 17 | Chihuahua | FS | 2.8 | 81 | CCD |
| 5 | 13 | Australian Shepherd | FS | 18.0 | 74 | CCD |
| 6 | 14 | Chihuahua | FS | 7.8 | 6 | CH |
| 7 | 13 | Chihuahua | FS | 3.3 | 6 | CH |
| 8 | 10 | Lab mix | FS | 34.5 | 4 | CH |
| 9 | 10 | German Pointer | FS | 31.0 | 2 | CH |
| 10 | 10 | Boxer | MN | 38.2 | 4 | CH |

CCD: canine cognitive dysfunction; CH: clinically healthy; FS: female spayed; MN: male neutered

**Table S2.**
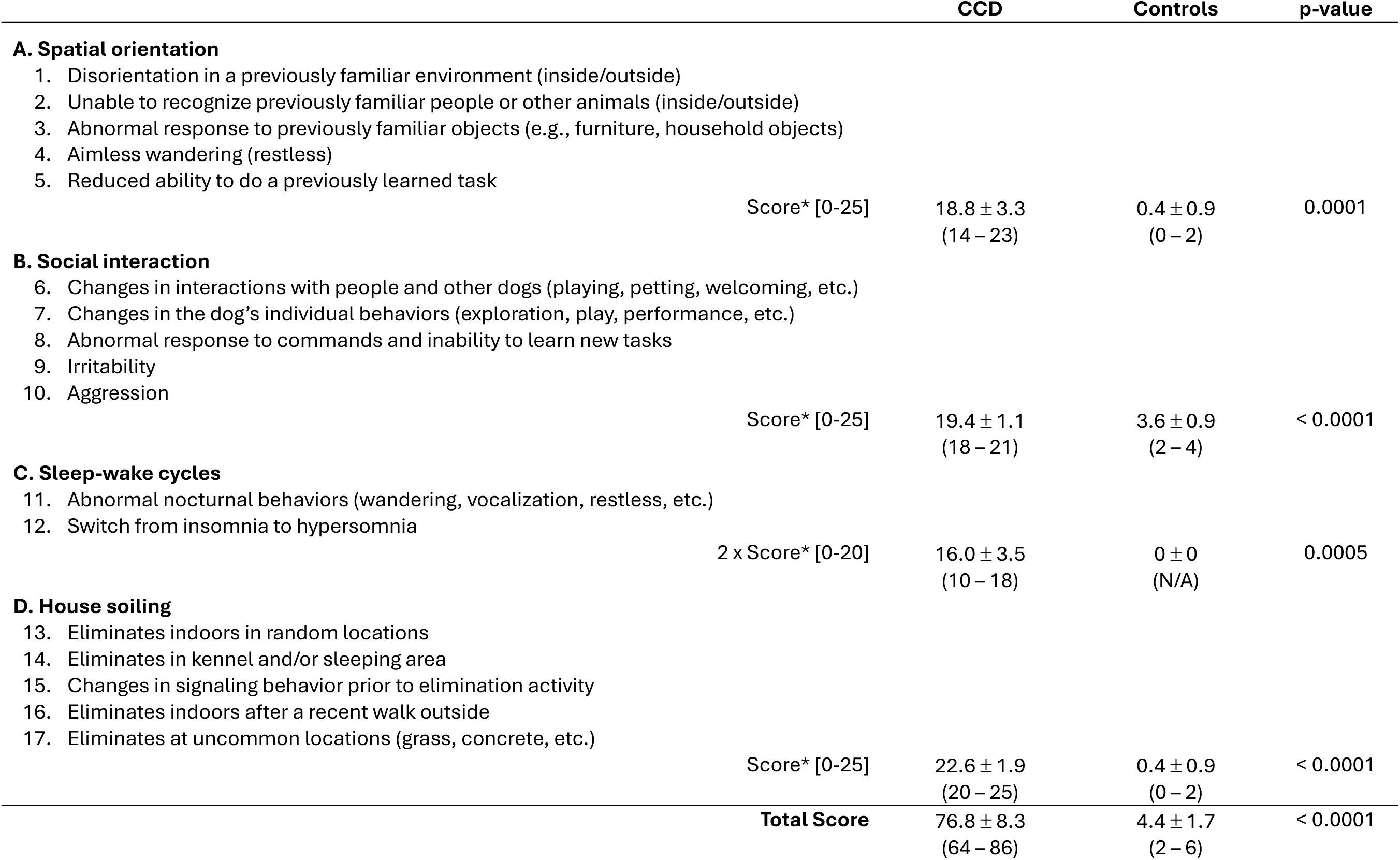
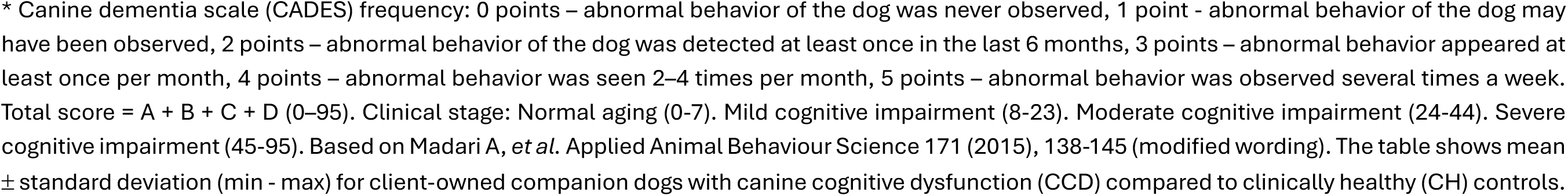
Mean canine dementia scale (CADES) scores in dogs with canine cognitive dysfunction (CCD) vs controls.

|  |  | CCD | Controls | p-value |
| --- | --- | --- | --- | --- |
| **A. Spatial orientation** |  |  |  |  |
| 1. Disorientation in a previously familiar environment (inside/outside) |  |  |  |  |
| 2. Unable to recognize previously familiar people or other animals (inside/outside) |  |  |  |  |
| 3. Abnormal response to previously familiar objects (e.g., furniture, household objects) |  |  |  |  |
| 4. Aimless wandering (restless) |  |  |  |  |
| 5. Reduced ability to do a previously learned task |  |  |  |  |
|  | Score* [0-25] | 18.8 ± 3.3<br>(14 – 23) | 0.4 ± 0.9<br>(0 – 2) | 0.0001 |
| **B. Social interaction** |  |  |  |  |
| 6. Changes in interactions with people and other dogs (playing, petting, welcoming, etc.) |  |  |  |  |
| 7. Changes in the dog's individual behaviors (exploration, play, performance, etc.) |  |  |  |  |
| 8. Abnormal response to commands and inability to learn new tasks |  |  |  |  |
| 9. Irritability |  |  |  |  |
| 10. Aggression |  |  |  |  |
|  | Score* [0-25] | 19.4 ± 1.1<br>(18 – 21) | 3.6 ± 0.9<br>(2 – 4) | < 0.0001 |
| **C. Sleep-wake cycles** |  |  |  |  |
| 11. Abnormal nocturnal behaviors (wandering, vocalization, restless, etc.) |  |  |  |  |
| 12. Switch from insomnia to hypersomnia |  |  |  |  |
|  | 2 x Score* [0-20] | 16.0 ± 3.5<br>(10 – 18) | 0 ± 0<br>(N/A) | 0.0005 |
| **D. House soiling** |  |  |  |  |
| 13. Eliminates indoors in random locations |  |  |  |  |
| 14. Eliminates in kennel and/or sleeping area |  |  |  |  |
| 15. Changes in signaling behavior prior to elimination activity |  |  |  |  |
| 16. Eliminates indoors after a recent walk outside |  |  |  |  |
| 17. Eliminates at uncommon locations (grass, concrete, etc.) |  |  |  |  |
|  | Score* [0-25] | 22.6 ± 1.9<br>(20 – 25) | 0.4 ± 0.9<br>(0 – 2) | < 0.0001 |
| **Total Score** |  |  |  |  |
|  |  | 76.8 ± 8.3<br>(64 – 86) | 4.4 ± 1.7<br>(2 – 6) | < 0.0001 |

\* Canine dementia scale (CADES) frequency: 0 points – abnormal behavior of the dog was never observed, 1 point - abnormal behavior of the dog may have been observed, 2 points – abnormal behavior of the dog was detected at least once in the last 6 months, 3 points – abnormal behavior appeared at least once per month, 4 points – abnormal behavior was seen 2–4 times per month, 5 points – abnormal behavior was observed several times a week. Total score = A + B + C + D (0–95). Clinical stage: Normal aging (0–7). Mild cognitive impairment (8–23). Moderate cognitive impairment (24–44). Severe cognitive impairment (45–95). Based on Madari A, *et al.* Applied Animal Behaviour Science 171 (2015), 138–145 (modified wording). The table shows mean ± standard deviation (min - max) for client-owned companion dogs with canine cognitive dysfunction (CCD) compared to clinically healthy (CH) controls.

**Table S3.** Hematological and serum biochemical profiles of dogs with canine cognitive dysfunction (CCD) vs controls.

| Complete Blood Count |  |  |  |  |  |  |  |  |  |
| --- | --- | --- | --- | --- | --- | --- | --- | --- | --- |
|  | CCD | Controls | p-value | Reference |  | CCD | Controls | p-value | Reference |
| Erythrocytes (M/μL) | 6.52 ± 0.73<br>(5.25 - 7.15) | 7.27 ± 1.26<br>(5.85 - 9.03) | 0.29 | 5.65 - 8.87 | Lymphocytes (K/μL) | 1.67 ± 0.77<br>(0.96 - 2.83) | 1.83 ± 0.44<br>(1.32 - 2.30) | 0.70 | 1.05 - 5.10 |
| Hematocrit (%) | 40.0 ± 3.8<br>(33.4 - 43.3) | 47.7 ± 7.7<br>(39.1 - 56.3) | 0.09 | 37.3 - 61.7 | Monocytes (K/μL) | 0.76 ± 0.49<br>(0.41 - 1.60) | 0.51 ± 0.05<br>(0.43 - 0.55) | 0.32 | 0.16 - 1.12 |
| Hemoglobin (g/dL) | 14.9 ± 1.5<br>(12.3 - 16.3) | 16.8 ± 3.0<br>(13.3 - 20.3) | 0.24 | 13.1 - 20.5 | Eosinophil (K/μL) | 0.67 ± 0.78<br>(0.24 - 2.07) | 0.31 ± 0.14<br>(0.09 - 0.44) | 0.35 | 0.06 - 1.23 |
| Leukocytes (K/μL) | 12.18 ± 8.68<br>(7.18 - 27.29) | 11.18 ± 4.11<br>(8.48 - 18.40) | 0.82 | 5.05 - 16.76 | Basophil (K/μL) | 0.05 ± 0.05<br>(0 - 0.11) | 0.04 ± 0.02<br>(0.02 - 0.07) | 0.72 | 0.00 - 0.10 |
| Neutrophils (K/μL) | 9.03 ± 7.57<br>(4.73 - 22.46) | 8.49 ± 3.86<br>(5.81 - 15.32) | 0.89 | 2.95 - 11.64 | Platelets (K/μL) | 415 ± 239<br>(117 - 763) | 345 ± 123<br>(185 - 524) | 0.58 | 148 - 484 |
| Serum Chemistry Panel |  |  |  |  |  |  |  |  |  |
|  | CCD | Controls | p-value | Reference |  | CCD | Controls | p-value | Reference |
| Glucose (mg/dL) | 104 ± 13<br>(85 - 120) | 110 ± 17<br>(88 - 134) | 0.55 | 74 - 143 | Chloride (mmol/L) | 115 ± 3<br>(112 - 118) | 113 ± 4<br>(108 - 118) | 0.45 | 109 - 122 |
| Creatinine (mg/dL) | 0.9 ± 0.4<br>(0.6 - 1.6) | 0.8 ± 0.2<br>(0.5 - 1.1) | 0.71 | 0.5 - 1.8 | Total Protein (g/dL) | 7.1 ± 0.3<br>(6.8 - 7.6) | 7.0 ± 0.4<br>(6.5 - 7.6) | 0.49 | 5.2 - 8.2 |
| BUN (mg/dL) | 22 ± 11<br>(11 - 40) | 16 ± 6<br>(9 - 25) | 0.27 | 7 - 27 | Albumin (g/dL) | 3.4 ± 0.3<br>(3.0 - 3.7) | 3.4 ± 0.4<br>(3.0 - 4.1) | 0.93 | 2.3 - 4.0 |
| Phosphorus (mg/dL) | 4.4 ± 1.0<br>(3.0 - 5.9) | 4.5 ± 1.2<br>(3.2 - 6.1) | 0.91 | 2.5 - 6.8 | ALT (U/L) | 74 ± 41<br>(31 - 127) | 48 ± 13<br>(27 - 60) | 0.24 | 10 - 125 |
| Calcium (mg/dL) | 10.3 ± 0.2<br>(10.1 - 10.7) | 9.9 ± 0.7<br>(9.3 - 11.0) | 0.32 | 7.9 - 12.0 | ALP (U/L) | 245 ± 174<br>(62 - 460) | 195 ± 157<br>(48 - 366) | 0.64 | 23 - 212 |
| Sodium (mmol/L) | 150 ± 2<br>(147 - 153) | 147 ± 1<br>(146 - 149) | 0.08 | 144 - 160 | GGT (U/L) | 0.6 ± 0.9<br>(0 - 2) | 0.6 ± 0.5<br>(0 - 1) | 1.00 | 0 - 11 |
| Potassium (mmol/L) | 5.0 ± 0.6<br>(4.3 - 5.8) | 4.5 ± 0.4<br>(4.2 - 5.0) | 0.20 | 3.5 - 5.8 | Bilirubin (mg/dL) | 0.2 ± 0.1<br>(0.1 - 0.4) | 0.3 ± 0.2<br>(0.1 - 0.6) | 0.56 | 0.0 - 0.9 |

Values reflect mean ± std (min – max) for dogs with canine cognitive dysfunction compared to controls. Reference values were provided by IDEXX.

